# CRONOSOJA: a daily time-step hierarchical model predicting soybean development across maturity groups in the Southern Cone

**DOI:** 10.1101/2023.09.18.558336

**Authors:** Alan D. Severini, Santiago Álvarez-Prado, María E. Otegui, Monika Kavanová, Claudia R. C. Vega, Sebastián Zuil, Sergio Ceretta, Martín Acreche, Fidencia Amarilla, Mariano Cicchino, María E. Fernández-Long, Aníbal Crespo, Román Serrago, Daniel J. Miralles

**Affiliations:** INTA. Centro Regional Buenos Aires Norte. Estación Experimental Agropecuaria Pergamino. Argentina; Universidad Nacional de Rosario. Argentina; Consejo Nacional de Investigaciones Científicas y Técnicas (CONICET), Argentina; Facultad de Agronomía, Universidad de Buenos Aires. Argentina; INIA La Estanzuela. Uruguay; INTA. Centro Regional Córdoba. Estación Experimental Agropecuaria Manfredi. Argentina; INTA. Centro Regional Santa Fe. Estación Experimental Agropecuaria Rafaela. Argentina; INTA. Centro Regional Salta Jujuy. Estación Experimental Agropecuaria Salta. Argentina; Centro de Investigación Capitán Miranda, Instituto Paraguayo de Tecnología Agraria, Itapúa, Paraguay; INTA. Centro Regional Buenos Aires Sur. Estación Experimental Cuenca del Salado. Argentina

## Abstract

Accurate prediction of phenology is the most critical aspect for the development of models aimed at estimating seed yield, particularly in species that exhibit variable sensitivity to environmental factors throughout the cycle and among genotypes. With this purpose, we evaluated the phenology of 34 soybean varieties in 43 field experiments located in Argentina, Uruguay, and Paraguay. Experiments covered a broad range of maturity groups (2.2-6.8), sowing dates (from spring to summer), and latitude range (24.9-35.6 °S), thus ensuring a wide range of thermo-photoperiodic conditions during the growing season. Based on the observed data, daily time-step models were developed and tested, first for each genotype, and then across maturity groups. We identified base temperatures specific for different developmental stages and an extra parameter for calculating the photoperiod effect after the R1 stage (flowering). Also, an optimum photoperiod length for each maturity group was found. Model selection showed that the determinants of phenology across maturity groups were mainly affecting the duration of vegetative and early reproductive stages. Even so, early phases of development were better predicted than later ones, particularly in locations with cool growing seasons, where the model tended to overestimate their duration. Overall, we developed a soybean phenology model that, owing to its process-based nature, is able to simulate phenology well across widespread locations and sowing dates yet, because of its simplicity, can be adopted by a wide audience. The final model was made available at http://cronosoja.agro.uba.ar.

## 1 Introduction

High efficiency in the use of resources to obtain the highest net margin per unit area, in an environmentally sustainable structure, is necessary in non-subsidised economies. This has led to farming systems relying on more intensive use of land and reduced tillage. In Argentina and Uruguay, double-cropping soybean after wheat or barley is a key component of these systems; but delaying soybean planting beyond the optimum date determine reductions in yield of soybean up to 50 kg ha^-1^ day^-1^ for high latitude environments (Calviño et al., 2003). Yield losses in soybean caused by delayed sowing date can be partially mitigated by changing the maturity group (MG) of the cultivar, thus changing the sensitivity to photoperiod and consequently the length and timing of developmental stages. These changes allow adjusting soybean phenology to each specific environmental condition, which is crucial for avoiding yield losses.

Crop adaptation based on phenology is important for facing stressful environments. For instance, if flowering is too early, plant growth may be insufficient to produce a minimum level of biomass compatible with reasonable levels of yield (Mayers et al., 1991). Therefore, in those conditions where the growing season is extremely short (e.g. soybean at late sowing dates or at relatively high latitudes for a tropical crop) early vigour of the crop is far more important than the use of “full season” crops, which may delay flowering and reduce the period available for grain filling (Lawn et al., 1995). The mentioned extremes define the length of the growing season, and pre-flowering development may be manipulated to improve adaptation through balancing the optimum time of flowering and the consequent duration of pre- and post-flowering phases.

The main environmental factors identified as drivers of soybean development are photoperiod and temperature (Cober et al., 2001). Soybean is described as a “quantitative short day plant” as the maximum rate of development is observed when plants are exposed to short-day conditions (i.e. below a threshold or optimum photoperiod). The rate of development is reduced when the photoperiod is greater than the optimum value (Summerfield, 1998). Photoperiod sensitivity and optimum photoperiod values show variations with the maturity group. Cultivars from low maturity groups present a lower sensitivity and a higher optimum photoperiod than genotypes of high maturity groups (Summerfield, 1998). Temperature per se is the other factor affecting the rate of development which is the only one that has a universal impact on the rates of development (Aitken, 1974), meaning that all crops and all phases of development are sensitive to temperature (Miralles and Slafer, 1999). In general, the higher the temperature the faster the rate of development and consequently the shorter the time to complete a particular developmental phase (Garner and Allard, 1930; Miralles and Slafer, 1999; Slafer and Rawson, 1994).

In soybean, as in many other crops, the number of grains per unit area is the yield component best correlated with yield variations (Egli, 1998). Although the number of grains per unit area is determined throughout the crop cycle, this yield component in soybean is largely determined by the events occurring soon after flowering up to early seed filling (i.e. from R3 to R6 according to the scale of Fehr and Caviness (1977); Monzon et al. (2021)). During that period, named “critical period for yield”, in the absence of nutrient and water restriction, variations in accumulated solar radiation during pod setting are closely associated with changes in the number of grains per unit area (Calviño et al., 2003; Kantolic and Slafer, 2001; Nico et al., 2015). Additionally, during that critical period the variations in the number of grains per unit area are mostly associated with variations in the number of pods per unit area (Jiang and Egli, 1995; Monzon et al., 2021; Nico et al., 2019). As overlapping between grain number and grain weight determination is common in soybean, the environmental conditions at which the critical period is exposed determine not only the number of grains but also grain weight and thereby final yield (Monzon et al., 2021). A positive relationship has been found between the duration of the critical period and yield (Dunphy et al., 1979), as well as between yield and the seed number per unit area (Egli and Bruening, 2000; Kantolic and Slafer, 2007). Therefore, delays in sowing date have a negative impact on yield because they generally expose the critical period to less favourable growing conditions, such as shorter days and reduced incident radiation as well as temperature (Egli and Bruening, 2000; Kantolic et al., 2013; Nico et al., 2015). At the light of evidences described above in terms of the importance of the environmental conditions during the “critical period” for soybean’s yield determination, accurate prediction of its occurrence is important for assisting farmers on the genotype and sowing date selection.

Simulation models like APSIM (Keating et al., 2003) and DSSAT (Jones et al., 2003) are extremely useful to predict crop phenology and seed yield under different environments. However, in these models the calibration of genetic coefficients for simulating phenology and yield is a complex and time-consuming process limiting the number of genotypes available for simulation. Consequently, the modelled genotypes may not be representative of the broad range of genetic material grown by farmers, limiting the scope of the tool for farmer management decisions. To tackle this issue in a simple way, simulation models use generic coefficients based on maturity groups and growth habits (Salmerón and Purcell, 2016).

Here we aimed at constructing a model, named CRONOSOJA, simple enough to be calibrated for many varieties available at the market, yet able to predict phenology with reasonable accuracy across Argentina, Paraguay and Uruguay. We aimed at (i) characterising the genotypic variability of phenology within the current commercial soybean varieties of Argentina, Uruguay and Paraguay, and (ii) building a simple dynamic model based on photothermal conditions of different regions to predict different phenological stages in soybean. The model outputs are currently available — in Spanish language — on an interactive website (http://cronosoja.agro.uba.ar/index.php/), which also includes estimates of frost risk and water availability during the growing season.

## 2 Materials and Methods

### 2.1 Sites, field experiments and genotypes

Forty-three field experiments were established in Argentina, Paraguay, and Uruguay, from 2016 to 2020 (Figure 1, Table 1). Each experiment consisted of 34 soybean varieties of different maturity groups (MG) from 2 to 6, representing some of the most frequently sown varieties in these countries (Table 2). In order to explore a wide range of temperature and photoperiod conditions, experiments were sown from the middle of October to the end of January, with a gap of around one month between successive sowing dates at different locations (Table 1). Weeds, insects, and diseases were controlled to ensure optimal growing conditions.

**Table 1:**
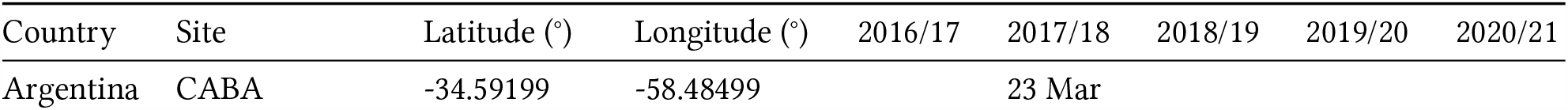
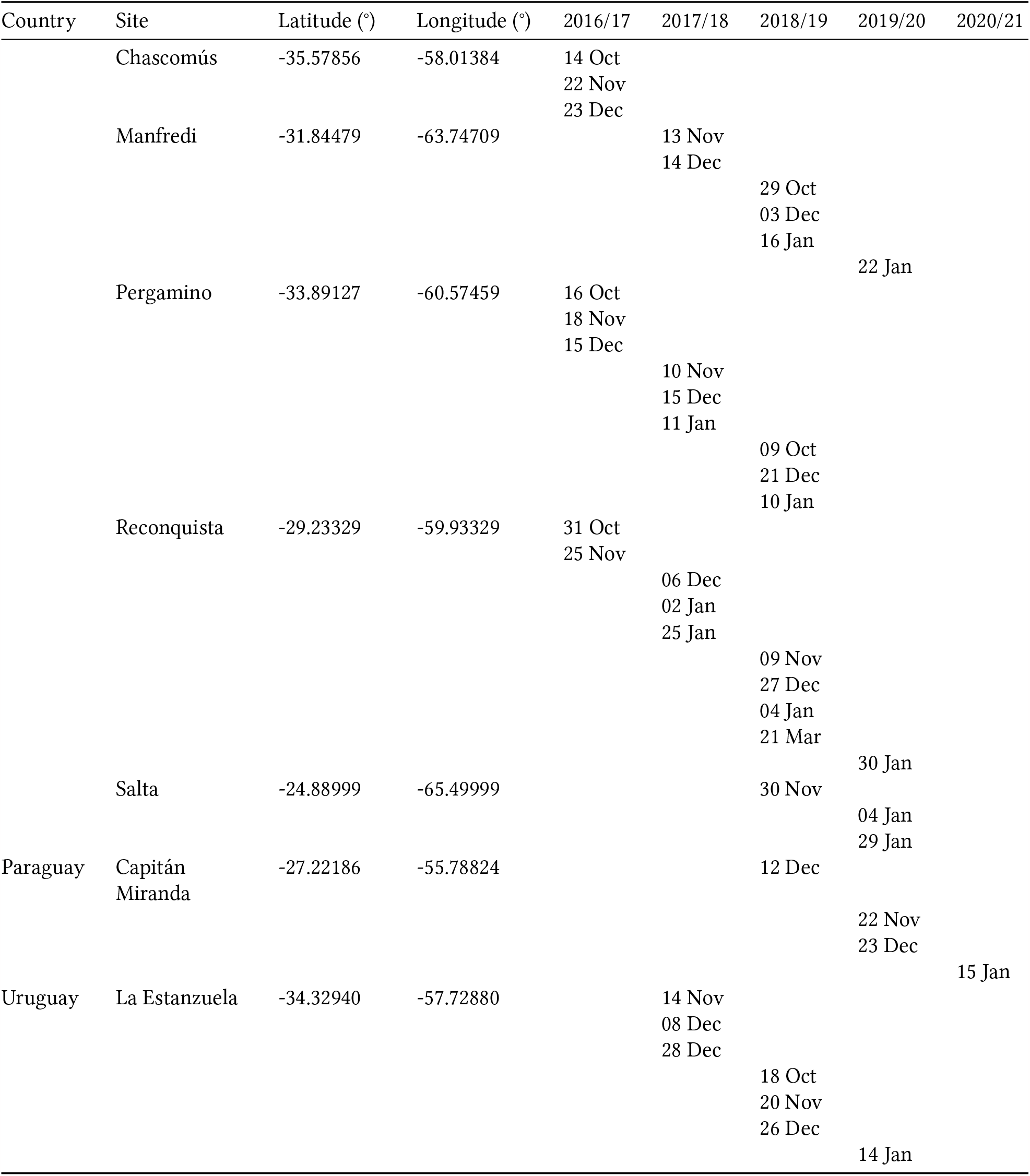
The location of experimental sites, and sowing dates tested in each of the four experimental seasons.

**Table 2:** Genotypes used in field experiments, including their maturity group (MG) and breeding company.

| MG | Company | Genotype |
| --- | --- | --- |
| 2.5 | GDM | DM 2200 |
| 2.5 | Nidera | NS 2632 |
| 3.0 | Nidera | NS 3220 |
| 3.5 | GDM | DM 3312 |
| 3.5 | Nidera | NS 3809 |
| 3.7 | Ferias del Norte | FN 3.85 |
| 3.9 | Santa Rosa Semillas | RA 349 |
| 4.0 | GDM | DM 3815 |
| 4.0 | GDM | DM 40R16 |
| 4.1 | Ferias del Norte | FN 4.35 |
| 4.2 | Asgrow | AW 4326 |
| 4.5 | GDM | DM 4612 |
| 4.7 | Bayer | CZ 4505 |
| 4.9 | Bayer | CZ 4.97 |
| 4.9 | Nidera | NS 4955 |
| 5.0 | GDM | DM 50i17 |
| 5.1 | Nidera | NS 5258 |
| 5.5 | GDM | DM 53i53 |
| 5.5 | INIA Uruguay | Génesis 5501 |
| 5.5 | INTA | INTA Paraná 5500 |
| 5.5 | Santa Rosa Semillas | RA 550 |
| 5.6 | INIA Uruguay | Génesis 5601 |
| 5.6 | INIA Uruguay | Génesis 5602 |
| 5.7 | Asgrow | AW 5714 |
| 5.7 | Asgrow | AW 5815 |
| 5.7 | Bayer | CZ 5905 |
| 5.9 | Santa Rosa Semillas | RA 549 |
| 6.0 | GDM | DM 6.2i |
| 6.2 | INIA Uruguay | Génesis 6201 |
| 6.2 | INTA | INTA Paraná 6200 |
| 6.3 | Bayer | CZ 6505 |
| 6.5 | Asgrow | AW M6410 |
| 6.5 | Nidera | NS 6248 |
| 6.8 | Santa Rosa Semillas | RA 655 |

**Figure 1:**
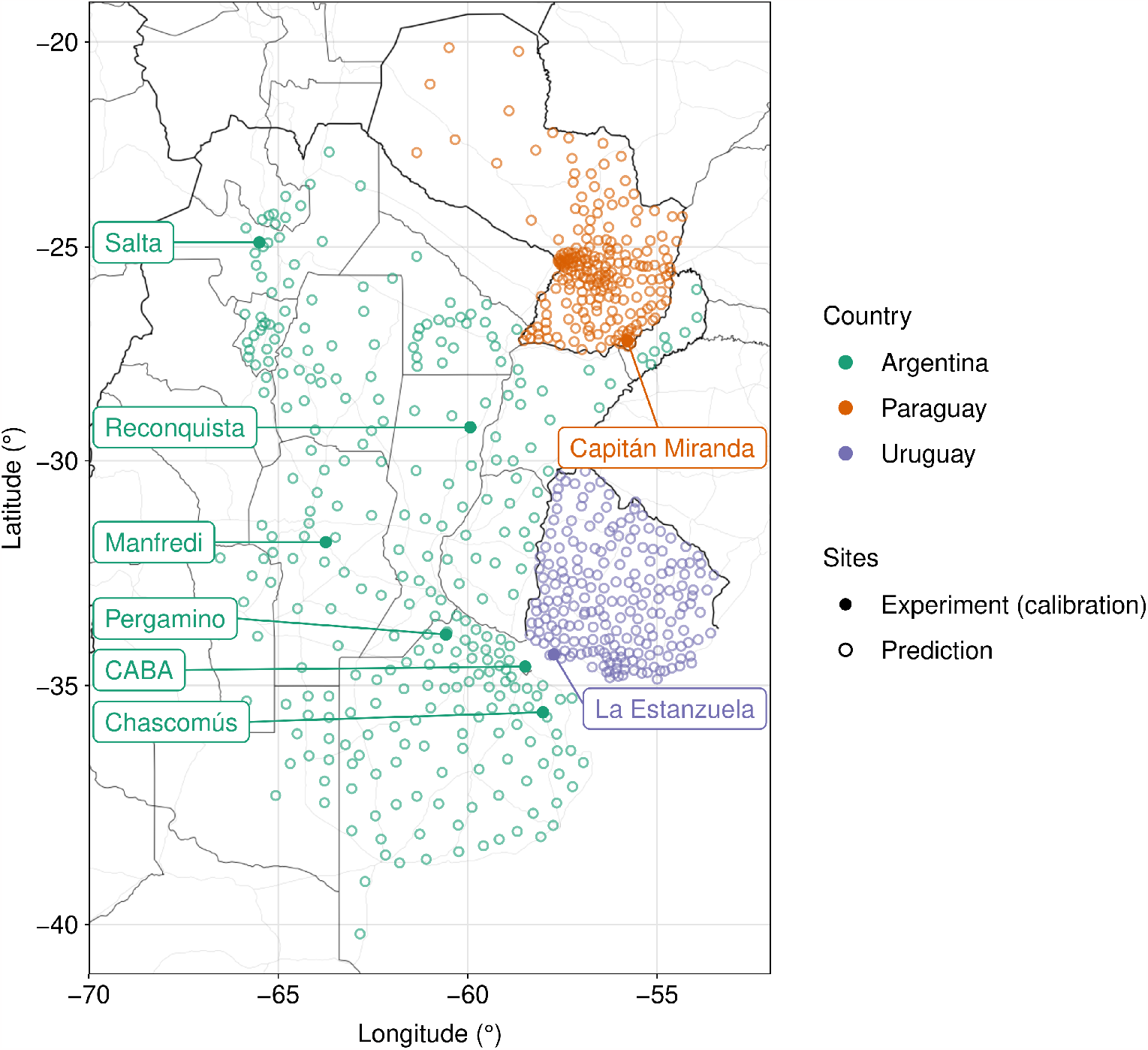
Distribution of experimental sites (solid symbols) and prediction sites (empty symbols) across Argentina, Paraguay and Uruguay.

At each experimental site, the 34 genotypes detailed in Table 2 were sown simultaneously in a wide range of sowing dates in field plots of 4 rows wide and 5 m long with a density of ca. 25 pl m^-2^ (early sowings) and ca. 30 pl m^-2^ (late sowings) under no water and nutritional restrictions. The experimental design was a randomised complete block design with three repetitions per genotype. Different phenological stages were recorded at least thrice weekly, plot by plot, by using the scale of Fehr and Caviness (1977) (see Table 3).

**Table 3:** Translation of the scale of Fehr and Caviness (1977) into a numeric variable (*R*).

| Stage | Abbreviation | $R$ |
| --- | --- | --- |
| Sowing | SO | 0 |
| Emergence | EM | 1 |
| End of juvenile | JU | 2 |
| Floral initiation | R0 | 3 |
| Flowering | R1 | 4 |
| Start pod | R3 | 5 |
| Start seed filling | R5 | 6 |
| Maximum seed size | R6 | 7 |
| Physiological maturity | R7 | 8 |
| Harvest maturity | R8 | 9 |

Environmental data collected included daily minimum and maximum temperature (°C) and photoperiod length (h). At the locations where weather stations were available, these data were recorded *in situ*, close to the experiments.At locations where weather stations were not available, or when there were temporal gaps in the data, we used data from Hersbach et al. (2018), downloaded from the Copernicus Climate Change Service (C3S) Climate Data Store. These gridded data were processed to extract temperature values from the exact coordinates of each site by means of bi-cubic interpolation with the software `CDO` (Schulzweida, 2021). Daily minimum and maximum temperatures were determined from hourly temperatures. Photoperiod was estimated given the site latitude and date with the package `geosphere` (Hijmans, 2021) in `R` (R Core Team, 2022).

### 2.2 Formulating a daily time-step developmental model

The phenological scale of Fehr and Caviness (1977) was translated into a numeric scale to be used as a regression dependent variable called *R* (Table 3).

We formulated a multiplicative model to calculate the development rate depending on daily photoperiod and temperature (Grimm et al., 1994; Major et al., 1975b). The equation describing the developmental stage (*R*) at a specified time (*t*, in days after sowing) at site *i* and genotype *j* was

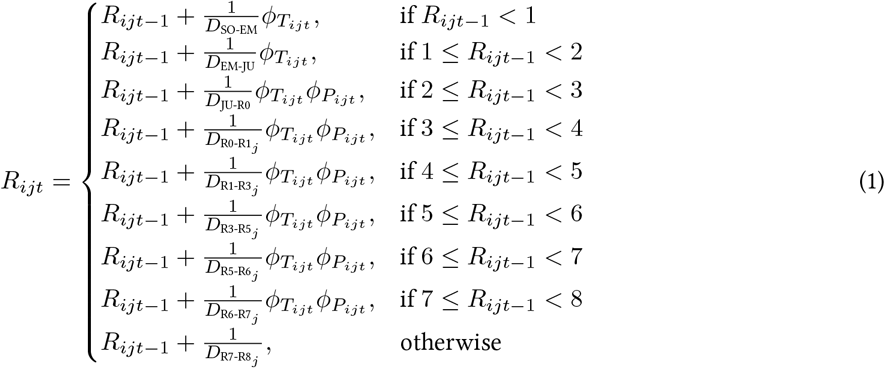

where *R*_*ijt−*1_ is the developmental stage from the previous day (*t* −1) at site *i* and genotype *j*; *D*_SO-EM_ and *D*_EM-JU_ are thermal days (i.e., equivalent to calendar days if temperature is optimum) for phases SO-EM and EM-JU, respectively; *D*_JU-R0_, *D*_R0-R1_, *D*_R1-R3_, *D*_R3-R5_, *D*_R5-R6_, *D*_R6-R7_, are photothermal days (i.e., equivalent to calendar days if temperature is optimum and photoperiod is less or equal to the optimum) for stages JU-R0, R0-R1, R1-R3, R3-R5, R5-R6, and R6-R7, respectively; and *D*_R7-R8_ are days for stage R7-R8. On the other hand, 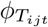and 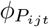are multipliers that describe the temperature and photoperiod responses, respectively, by adopting values between 0 and 1. Therefore, 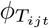and 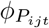penalise the maximum developmental rates (e.g. 1*/D*_R1-R3_). 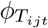and 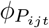are defined for site *i*, genotype *j* and day *t*, that is, these functions depend on the site because of its input variables and depend on the genotype because of its genotype-specific parameters.

For simulating the response of 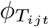to temperature we used the function of Wang and Engel (1998),

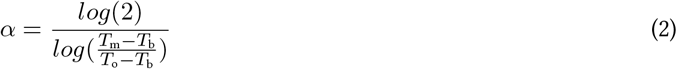

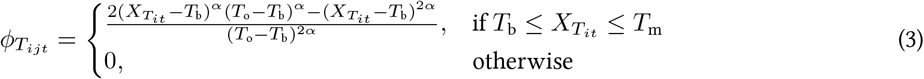

where 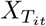is daily air temperature of site *i* and day *t, T*_b_ is the base temperature, *T*_o_ is the optimum temperature, and *T*_m_ is the maximum temperature, all in °C.

At this stage, the first alternative models were evaluated: one where *T*_b_ was constant across developmental phases, and another one where it differed between developmental stage ranges, so that

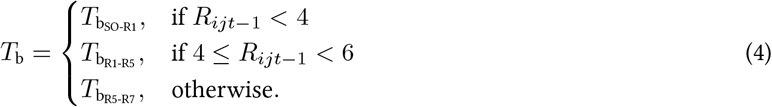

The temperature function was calculated sub-daily, at 3-hour intervals, by interpolating between daily minimum and daily maximum temperatures (Jones et al., 1986) using a cubic function as employed in APSIM (Keating et al., 2003). The resulting eight, sub-daily temperature multipliers were averaged to represent a daily temperature multiplier 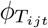.

We wanted to test competing models at this stage, since current models differ in their structure. For instance, DSSAT includes a parameter named `R1PPO`, which increases photoperiod sensitivity after R1 by reducing the optimum photoperiod by an amount that depends on the genotype. APSIM, however, lacks this parameter. Thus, we tested the alternative of including or excludingΔ*P*_o_, leading to four alternative models (Table 5).

For formalising the photoperiod multiplier, 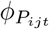, we formulated the second set of alternative equations: one where the photoperiod response was constant across reproductive development, and another one where the response threshold was reduced by the parameter 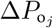(similar to parameter `R1PPO` in CROPGRO (Boote et al., 1998b)), making plants more sensitive to photoperiod after R1. Therefore, the simpler model for the photoperiod multiplier was

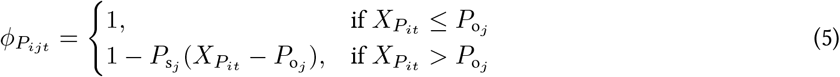

whereas the more complex one was

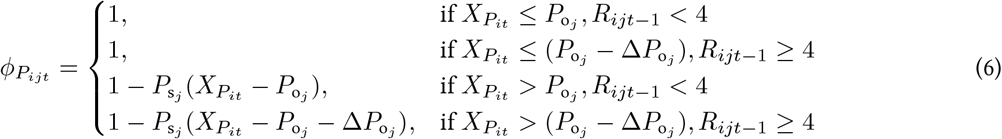

where 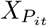is the photoperiod (h) of site *i* and day *t*, 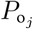is the optimum photoperiod of genotype *j*, 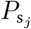is the photoperiod sensitivity of genotype *j*, and 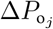represents the shift in the optimum photoperiod after R1 of genotype *j*.

### 2.3 Explaining initial variability in phenology

To understand the sources of variation affecting the duration of each developmental stage, we fitted a mixed effects regression explaining the duration of stages as a function of sowing date (days since 1st October of the current year), maturity group (as factor), genotype, site and year. Fixed effects were the effect of sowing date, maturity group and, when significant, their interaction. Random effects (random intercepts) were genotype, site and year. Models were fitted with package `lme4` (Bates et al., 2015) in `R`. Models with and without an interaction between sowing date and maturity group were compared by means of Akaike information criteria (AIC). The resulting best models included the interaction term for stages EM-R1, R1-R3, R3-R5, and R5-R6 (Equation 7), but lacked this term for stages R6-R7 and R7-R8 (Equation 8):

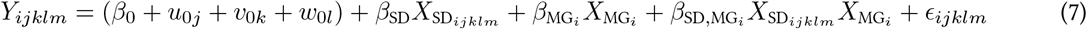

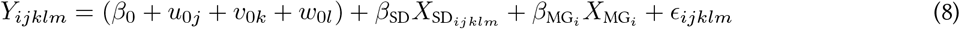

where *Y*_*ijklm*_ is the duration of the stage for maturity group *i*, genotype *j*, site *k*, year *l* and replicate *m*; *β*_0_ is the general intercept; *β*_SD_ is the slope against sowing date; 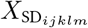is the sowing date (as days since 1st October) of maturity group *i*, genotype *j*, site *k*, year *l* and replicate 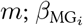is the effect of maturity group 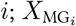is maturity group 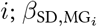is the effect of the interaction between sowing date and maturity group *i*; *u*_0*j*_ is the random effect of genotype *j*; *v*_0*k*_ is the random effect of site *k*; *w*_0*l*_ is the random effect of year *l*; and *ϵ*_*ijklm*_ is the error of maturity group *i*, genotype *j*, site *k*, year *l* and replicate *m*.

Once the appropriate models were fitted, we obtained the variance of each source of variation with the function `get_variances` from the `R` package `insight` (Lüdecke et al., 2019). Since this resulted in a combined fixed effects variance (i.e. sowing date and maturity group variances were amalgamated, rendering them indistinguishable from one another), we further partitioned the fixed effects variance per source of variation using the `R` package `partR2` (Stoffel et al., 2020).

### 2.4 Fitting one model per genotype (genotype-based model)

The first step of the calibration was performed in a genotype-based manner. Parameters to be calibrated were photothermal days *D*_R0-R1_, *D*_R1-R3_, *D*_R3-R5_, *D*_R5-R6_, *D*_R6-R7_, calendar days *D*_R7-R8_, and those parameters contained in the temperature function 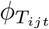and in the photoperiod response function 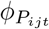. From 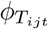, we chose to calibrate the base temperature *T*_b_, as judging from the literature was the parameter that changed the most (Jones et al., 1986; Setiyono et al., 2007). Parameters *T*_o_ and *T*_m_ were set to 30 and 45 °C, respectively. From 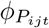, we calibrated all three parameters, that is, *P*_o_, *P*_s_, and, where present,Δ*P*_o_.

To perform the calibration, we coded a likelihood function in the hybrid language `Rcpp` (Eddelbuettel and François, 2011). This function ran the model per genotype, across all sowing dates and locations where the corresponding genotype was sown, iterating through possible solutions of *D*_R0-R1_, *D*_R1-R3_, *D*_R3-R5_, *D*_R5-R6_, *D*_R6-R7_, *D*_R7-R8_, *T*_b_, *P*_o_, *P*_s_, andΔ*P*_o_. The function returned a log-likelihood value that was maximised by means of a Metropolis-Hastings Markov Chain Monte Carlo (MCMC) implemented in the `R` package `BayesianTools` (Hartig et al., 2019). The MCMC was performed in four independent chains with 3 × 10^5^ iterations, which ensured that the chains mixed appropriately. Convergence was assured by the diagnostic 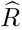, which was always less than 1.01 (Vehtari et al., 2021).

The prior distributions of most parameters were constructed by sampling from the literature, with the following modifications. First, their standard deviation was tripled to allow the model to explore other possible values given our different model structure. Second, in the case of *T*_b_, as we were implementing a different function than those used in Jones et al. (1986), Archontoulis et al. (2014) or Setiyono et al. (2007), we set them to have mean zero and a large standard deviation, since the actual values were unknown. Third, in the case of *D*_R3-R5_, their mean was calculated by subtracting from DSSAT parameter `FL.SD` (photothermal days from R1 to R5, Jones et al. (1986)) the parameter `FL.SH`, the latter being equivalent to *D*_R1-R3_. All priors were specified as truncated normal distributions with function `createTruncatedNormalPrior` from `BayesianTools`. Their final means, standard deviations, lower and upper limits are shown in Table 4. No priors were constructed for parameters *D*_SO-EM_, *D*_EM-JU_ and *D*_JU-R0_, which were given values 5, 5, and 6 thermal days, respectively, after consulting parameters from APSIM (presented by Archontoulis et al. (2014)).

**Table 4:** Mean, standard deviation, lower and upper limits for the truncated normal priors of each genotype-based model parameter.

| Parameter | Mean | Standard deviation | Lower limit | Upper limit |
| --- | --- | --- | --- | --- |
| $T_{bSO-R1}$ (°C) | 0.00 | 3.00 | -1000.0 | 10.0 |
| $T_{bR1-R5}$ (°C) | 0.00 | 3.00 | -1000.0 | 10.0 |
| $T_{bR5-R7}$ (°C) | 0.00 | 30.00 | -1000.0 | 10.0 |
| $D_{R0-R1}$ (d) | 6.00 | 9.00 | 1.0 | 30.0 |
| $D_{R1-R3}$ (d) | 7.50 | 4.00 | 1.0 | 30.0 |
| $D_{R3-R5}$ (d) | 7.63 | 2.00 | 1.0 | 30.0 |
| $D_{R5-R6}$ (d) | 13.70 | 10.00 | 1.0 | 40.0 |
| $D_{R6-R7}$ (d) | 13.70 | 10.00 | 1.0 | 40.0 |
| $D_{R7-R8}$ (d) | 12.00 | 3.00 | 1.0 | 30.0 |
| $P_o$ (h) | 13.30 | 1.00 | 10.0 | 16.0 |
| $P_s$ (h<sup>-1</sup>) | 0.29 | 0.03 | 0.0 | 0.6 |
| $\Delta P_o$ (h) | 0.00 | 0.40 | -1.0 | 1.0 |

**Table 5:** Univariate models tested by means of Pseudo-Bayesian model averaging (PBMA) and stacking (Yao et al., 2017). The higher the weight the better the model.

| Model | Temperature response | Photoperiod response after R1 | PBMA | Stacking |
| --- | --- | --- | --- | --- |
| u1 | $f(X_T, T_b)$ | $f(X_P, P_o, P_s)$ | 0.035 | 0.0 |
| u2 | | $f(X_P, P_o, P_s, \Delta P_o)$ | 0.073 | 0.0 |
| u3 | $f(X_T, T_{bSO-R1}, T_{bR1-R5}, T_{bR5-R7})$ | $f(X_P, P_o, P_s)$ | 0.273 | 0.0 |
| u4 | | $f(X_P, P_o, P_s, \Delta P_o)$ | 0.619 | 1.0 |

For model selection, we followed a multi-model inference approach to select the best models based on information criteria (Garibaldi et al., 2017). In each case, the most parsimonious model was chosen based on information criteria inferred by approximate leave-one-out cross-validation (LOO) with package `loo` (Vehtari et al., 2019) for `R`, which implements a fast computation of LOO using Pareto smoothed importance-sampling (PSIS-LOO, Vehtari et al. (2017)). For this, a second `Rcpp` function was coded that instead of yielding a cumulative log-likelihood resulted in a pointwise log-likelihood, i.e. a log-likelihood matrix with rows equal to the number of observations and columns equal to the number of iterations. The number of iterations was obtained from concatenating the four chains, discarding the burn-in fraction, and thinning every 60 iterations. The appropriate values of thinning were found after performing an autocorrelation test with the `R` package `coda` (Plummer et al., 2006). This point-wise log-likelihood was fed into the function `loo` from the package of the same name (Vehtari et al., 2019). Leave-one-out cross-validation, as implemented in the package `loo`, allowed us to calculate an expected log point-wise predictive density (ELPD). This per-genotype ELPD was concatenated into one vector (representing all genotypes) per each of the four competing models so that a combined ELPD per model was obtained. With these ELPDs, models were compared both by pseudo-Bayesian model averaging and by stacking (Yao et al., 2017), obtaining a probability (weight) per model. The model with the highest weight is the best model, that is, the one with best predictive accuracy and likely the one that fits the data best with the least amount of parameters. These are shown in Table 5.

### 2.5 Generating a common hierarchical model across genotypes

At the second stage of the calibration, we wanted to unify all 34 genotype-based models into one hierarchical model across maturity groups and genotypes. To do so we constructed a Bayesian multivariate regression that simultaneously predicted the parameters of the aforementioned best model using maturity group (MG) as a covariate. The model was multivariate in the sense that parameters *D*_R0-R1_, *D*_R1-R3_, *D*_R3-R5_, *D*_R5-R6_, *D*_R6-R7_, *D*_R7-R8_, *T*_b_, *P*_o_, *P*_s_, andΔ*P*_o_, were regressed simultaneously. The model was in turn a mixed-effects linear model, with MG as a fixed effect and genotype as a random effect (random intercepts). This model can be represented as

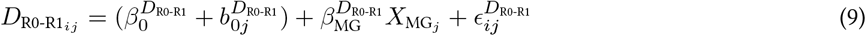

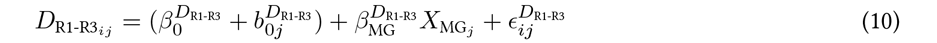

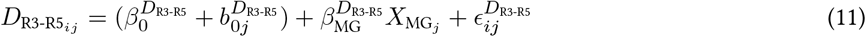

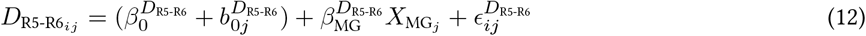

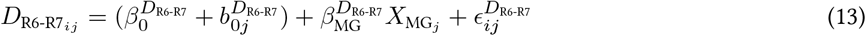

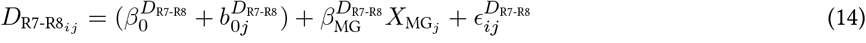

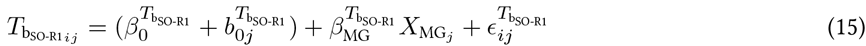

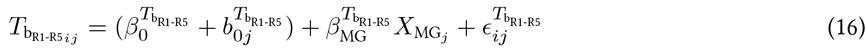

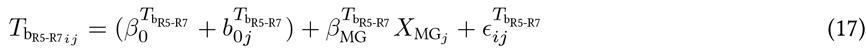

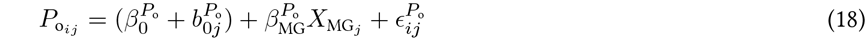

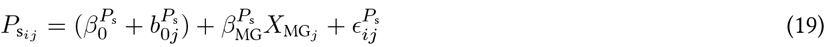

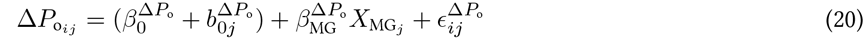

where 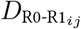is the *i* draw from parameter *D*_R0-R1_ of genotype 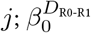is the global intercept and 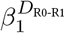the slope of the regression against 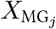, which represents maturity group from genotype *j*; and 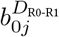and 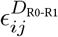are random variables representing, respectively, the random intercept of genotype *j*, and the random error from draw *i* and genotype *j*. The same nomenclature applies to the rest of the response variables.

Random effects of genotypes and errors were drawn from multivariate normal distributions (MVN) with mean 0 and variance-covariance matrices **Σ**_*b*_ and **Σ**_*ϵ*_. In the case of random intercepts resulting from genotypes, *b*_0*j*_, these were

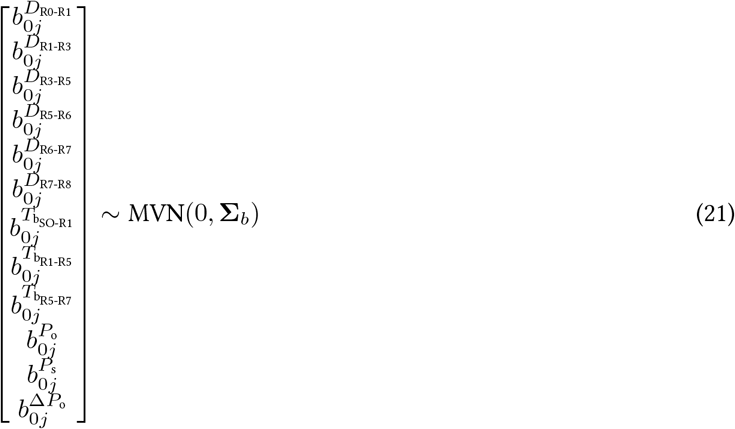

where 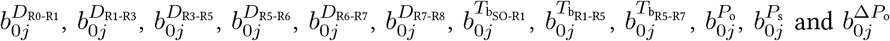and 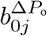are the random effects of genotype *j* over parameters *D*_R0-R1_, *D*_R1-R3_, *D*_R3-R5_, *D*_R5-R6_, *D*_R6-R7_, *D*_R7-R8_, 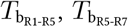, *P*_o_, *P*_s_ andΔ*P*_o_, respectively, and **Σ**_*b*_ is their variance-covariance matrix, with diagonal elements 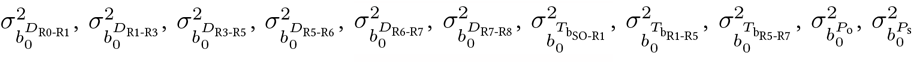and 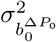showing the genotype variances and off-diagonal elements showing their covariances between different parameters (Section 9, Equation 36).

Errors *ϵ*_*ij*_ are correlated between parameters as well, so that

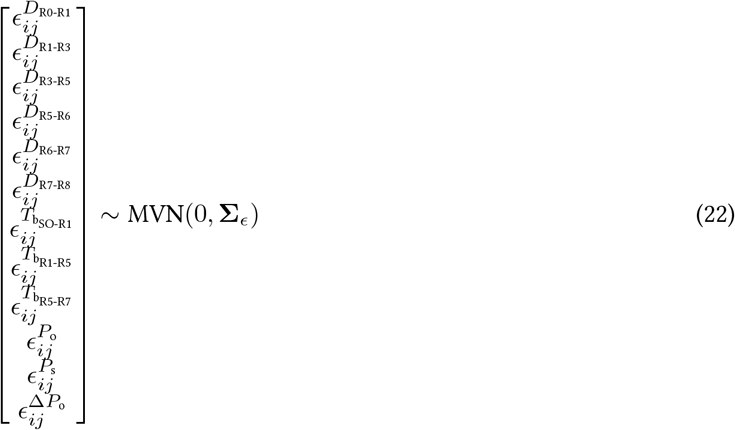

where **Σ**_*ϵ*_ is their variance covariance matrix, with its diagonal elements capturing the variances of the residuals of each response variable, and its off-diagonal elements representing the covariances between residuals of different response variables (Section 9, Equation 37).

During this stage of the calibration, the multivariate models were fitted with the probabilistic programming language `Stan` (Carpenter et al., 2017), accessed through package `brms` (Bürkner, 2018) in `R`. For fitting these models, theMCMC was run in 4 chains, where each chain had 2000 iterations, and the first 1000 were removed as part of the warm-up process, resulting in an MCMC sample of 4000 values. We used the default priors defined in the function `brm` from package `brms` (see Severini et al. (2020) for a detailed description). Convergence of the posterior density was evaluated again by the 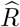test statistic, where values less than 1.01 indicated that the MCMC sampling process was adequate (Vehtari et al., 2021).

Given that ten model parameters were regressed simultaneously, there were competing models at this stage too, where parameters were allowed to be a function, or not, of MG. Several alternative models, possessing different parameters’ response to MG, were compared again using LOO by obtaining a probability (weight) per model (Vehtari et al., 2019).

Fixed and random effects are reported always as the median and its 90% credible-interval bounds (CI: lower limit, upper limit) of the posterior distribution.

### 2.6 Calibration and validation data sets

In order to have data both for calibrating the model as well as for testing it, we split the experimental data randomly within each genotype (i.e. a single genotype could contain multiple combinations of locations and sowing dates) into calibration (75%) and validation (25%) data sets with the function `initial_split` from the `R` package `rsample` (Silge et al., 2021). All subsequent calibrations, both genotype by genotype as well as the unified, hierarchical model, were done with the calibration data set. Data from the testing split were used to compare observed days after sowing (DAS) to R1, R3, R5, R6, R7 and R8 with the simulated DAS of each stage. This comparison was made by means of root mean squared error (RMSE), relative RMSE (RRMSE) and relative lack of accuracy (RLA) using the `R` package `metrica` (Correndo et al., 2021) from R. RLA measures the percentage of error given by how much a symmetric regression line (SMA) departs from the 1:1 line, representing systematic error (Correndo et al., 2021), in contrast with the relative lack of precision representing unsystematic error which can be visualised as random error around the SMA.

### 2.7 Predictions for website deployment

After calibrating the model and validating it against the validation dataset, the last step a was to generate predictions that served as inputs of the website. First, the main soybean producing regions of each country were identified after consulting local government public databases. Then, shapefiles for each country were downloaded from https://gadm.org/download_country_v3.html and one coordinate per province-district combination was calculated as a centroid of its shapefile polygon with package `sf` (Pebesma, 2018) in `R` (Figure 1). For every centroid coordinate, daily maximum and minimum temperatures for the last 10 years as well as daily photoperiod were determined in the same way as the calibration weather data processing detailed in Section 2.1. Sowing dates were simulated for every day from the beginning of October to the end of January for 10 years. For every sowing date day-of-year, genotype and location, days after sowing were averaged across 10 years for stages R1, R3, R5, R6, R7 and R8. These averages were later saved as CSV spreadsheets on a prediction-site basis to feed the website.

### 2.8 Available soil water estimation

To describe the temporal dynamics of available soil water, we used a soil water balance simulation model BHOA (for its acronym in Spanish: “Balance Hidrológico Operativo para el Agro”; Fernández-Long et al. (2013)). The BHOA is a one-layer bucket model that establishes a balance a balance between the atmospheric demand of water (given by the potential evapotranspiration; PET), the supply of water (given by the precipitation; PP) and the water stored in the ground (soil moisture; SM)

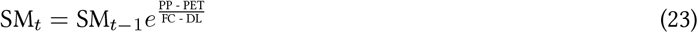

where SM_*t*_ is the soil moisture on day *t*, SM_*t−*1_ is the soil moisture on the day before, FC is the field capacity and DL is the drying limit, calculated as a percentage of the permanent wilting point (PWP). The available soil water is the difference between the amount of SM_*t*_ and the amount of PWP. This model has been proven to have a very good fit with observed data (Fernández-Long et al., 2018; Spennemann et al., 2020) and has been used in Argentina for the estimation of soil moisture with different objectives (Fernández-Long et al., 2021; Penalba et al., 2019; Peretti et al., 2023; Pinto et al., 2017).

## 3 Results

### 3.1 Environmental conditions explored across sites

The experiments allowed us to explore a wide range of temperatures and photoperiod combinations. From all the locations, Salta site was cooler when compared to the rest of the sites (explored temperatures within the growing season were between 6 and 30 °C) due to its higher altitude above sea level. Capitán Miranda site offered the most inductive conditions for soybean development across sites, i.e. the highest average temperatures (10 to 39 °C) and shortest photoperiods during the growing season (11.1 to 13.9 h, Figure 2).

**Figure 2:**
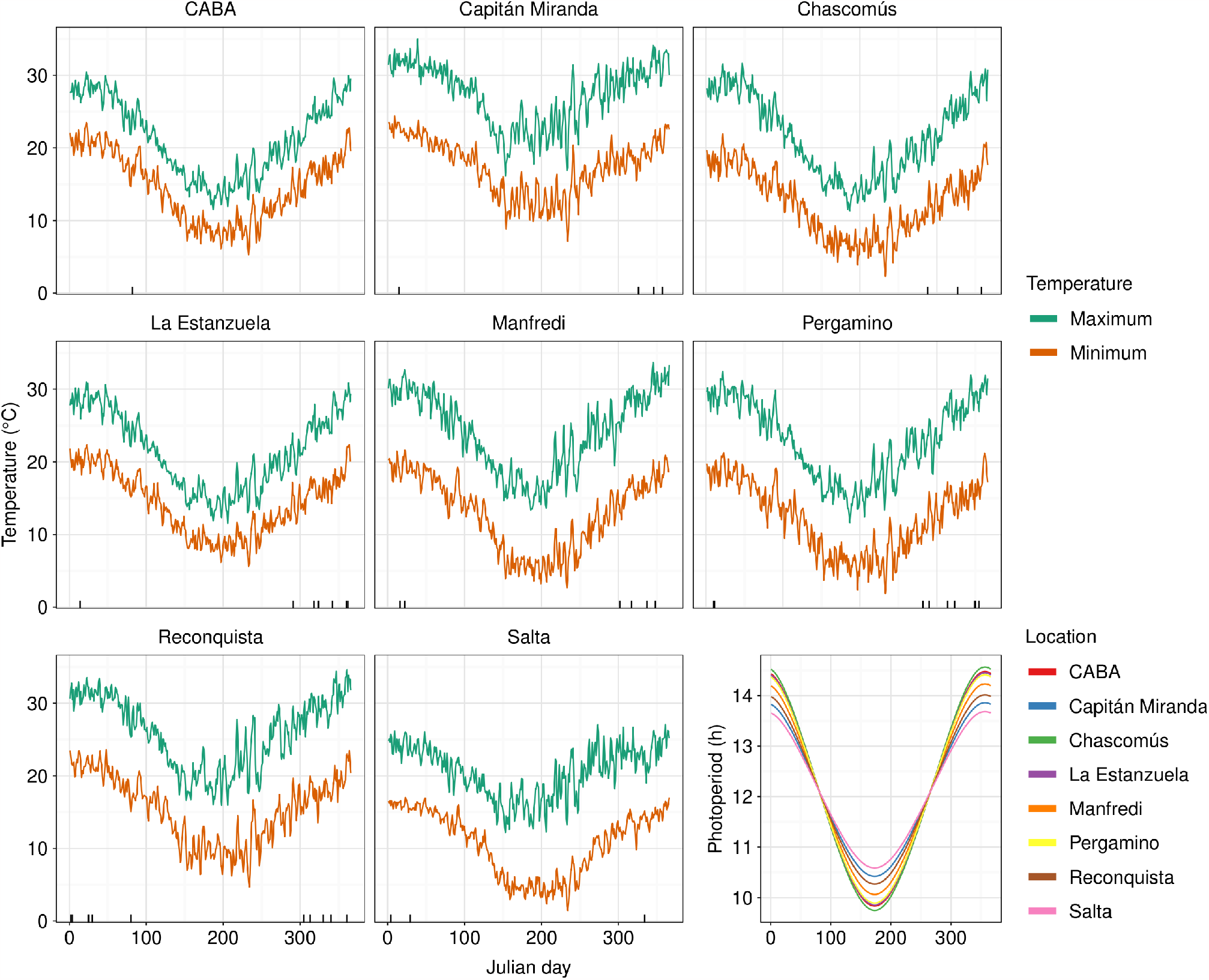
Average minimum and maximum daily temperatures for each experimental site during the evaluated years, and daily photoperiod for each location. Short, vertical segments along the x axis show the sowing dates at each location. The last sub-figure shows a summary of the photoperiod dynamics for all locations.

### 3.2 Genotypic and environmental variability in phenology

Due to the wide maturity group range of the evaluated soybean varieties (Table 2), site location (Table 1), and sowing dates from October to January across sites, a wide variation in days from sowing to developmental stages R1, R3, R5, R6, R7 and R8 (Fehr and Caviness, 1977) per genotype was observed (Figure 3). Variation in stages duration across genotypes and environments was reduced with advance of development, with coefficients of variation of 30.2, 25.1, 23.1, 20.7, 19.2, and 16.4% for stages R1, R3, R5, R6, R7, and R8, respectively. According to the phenological stage, this variation in the duration of the phases was mainly associated with the site and year (for all developmental stages). Sowing date and genotype affected all stages duration, with a smaller effects on R5-R6 and R7-R8 (Figure 4).

**Figure 3:**
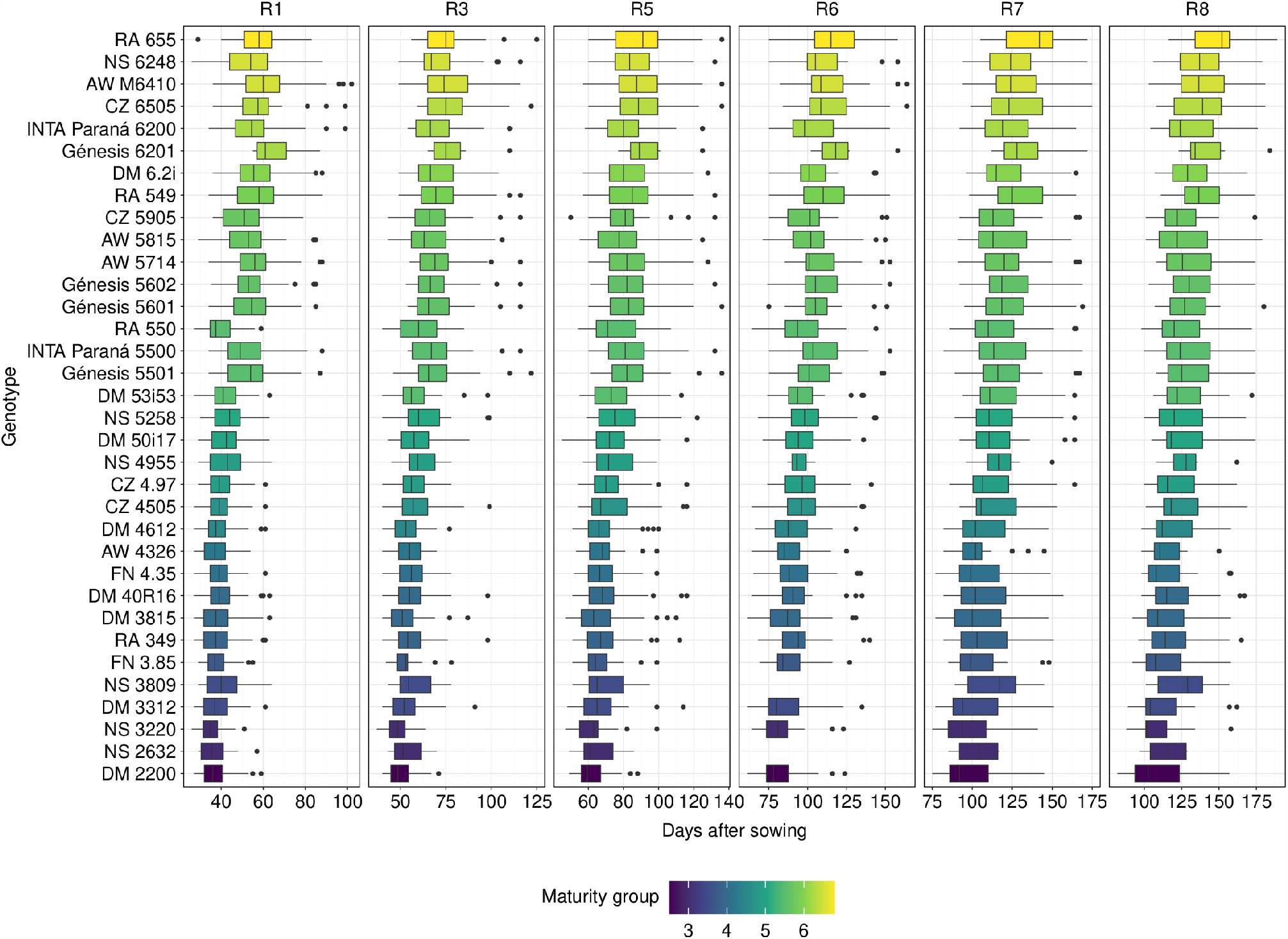
Variability in the developmental stages expressed as days from sowing to stages R1, R3, R5, R6, R7 and R8 (Fehr and Caviness, 1977). Each boxplot shows the data range explored across the sowing dates and locations as listed in Table 1.

**Figure 4:**
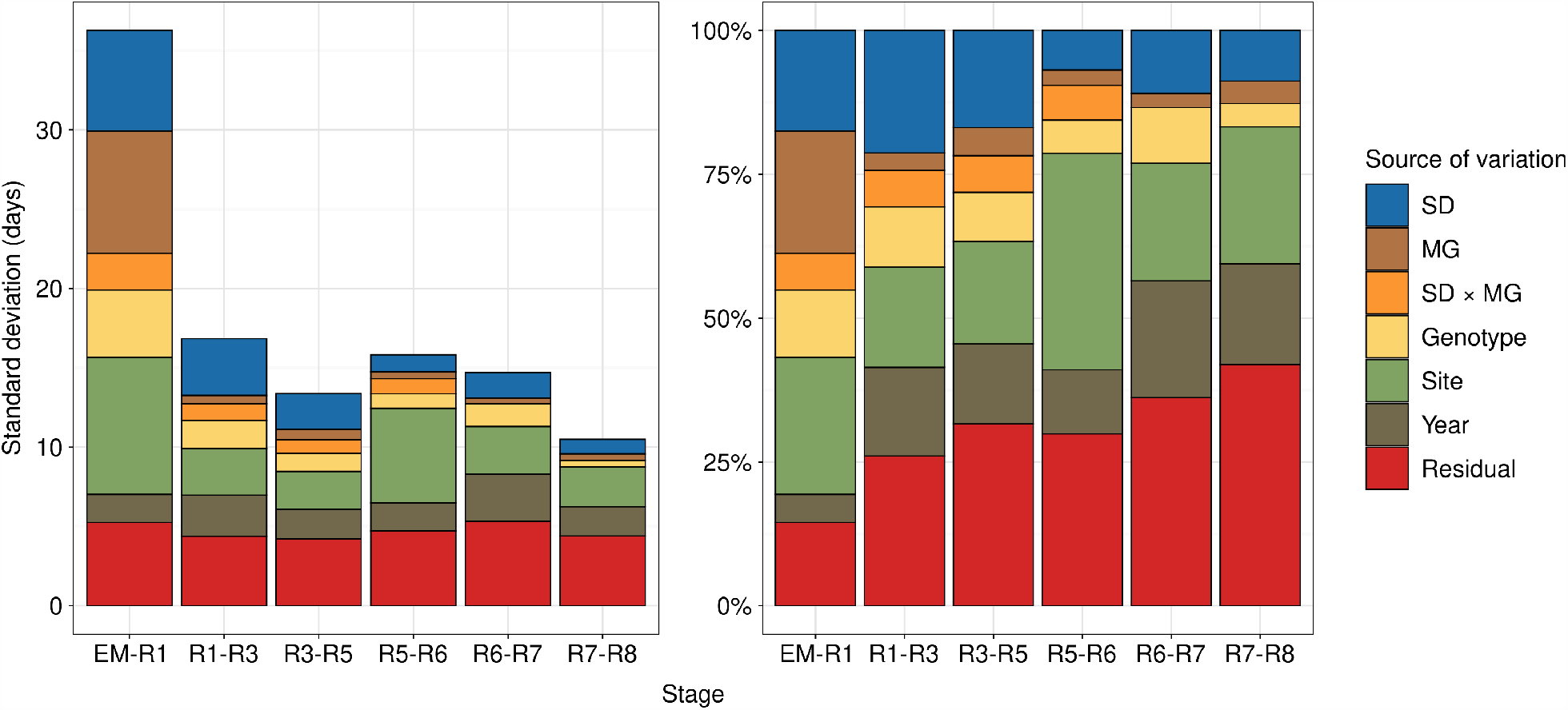
Standard deviation in stages duration partitioned per source of variation, represented as absolute (days, left) or relative (%, right) duration. Colours represent the different sources of variance such as sowing date (SD), maturity group (MG), and their interaction (SD × MG).

The EM-R1 phase was the one that showed the greatest variation in its duration and also was the one most affected by the MG and the genotype. Additionally, the genotypic variance given by both MG and genotype was decreasing with development, being the largest at the EM-R1 phase. Sowing date also produced larger variance early in the development, diminishing towards the end of the crop’s cycle. The variance attributed to site was the largest at EM-R1 and R5-R6 phases, while the variance associated with the year was constant across developmental stages when considering absolute values but increasing in relative terms as development progressed, behaving in the same manner as the residual error (Figure 4).

### 3.3 Parameters linked to developmental responses to the environment

To derive a unique hierarchical model, two model selection stages were incorporated. At the first selection stage, which chose the best univariate model on a genotype basis, the best model in terms of out-of-sample predictive accuracy was model `u4` (Table 5). Model `u4` had three base temperatures that differed across developmental stages 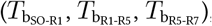, and an extra parameterΔ*P*_o_ for calculating the photoperiod multiplier after R1 (Table 5).

At the second calibration stage, which tested whether parameters from model `u4` were a function of maturity group, genotype, both or none, the best model was `m4` (Table 6), whose full notation is

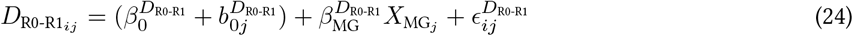

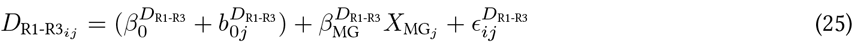

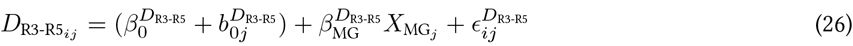

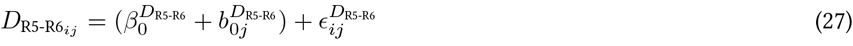

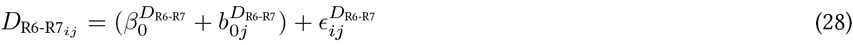

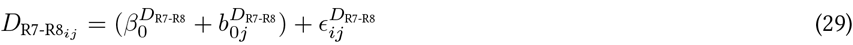

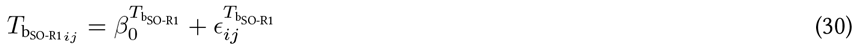

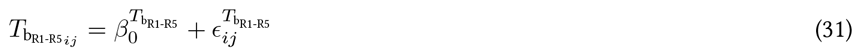

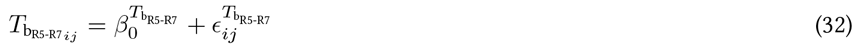

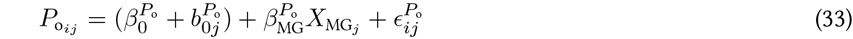

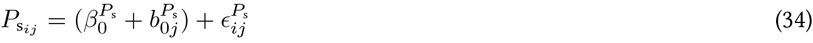

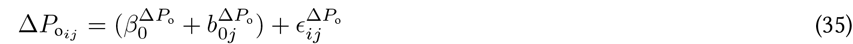

Parameters that concerned early plant development, i.e. vegetative and close to flowering stages, were the most affected by maturity group. Base temperatures did not change across maturity group or genotypes.

**Table 6:** Multivariate models differing in whether they included the terms 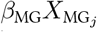and *b*_0*j*_, which respectively represent the linear response of the parameter to maturity group and the random intercept given by genotype. Their presence (1) or absence (0) is indicated separated by a slash, e.g. 1/1 indicates that the corresponding model included both 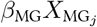and *b*_0*j*_. Models were compared by Pseudo-Bayesian model averaging (PBMA) and stacking (Yao et al., 2017). The higher the weight the better the model.

| Model | $D_{R0-R1}$ | $D_{R1-R3}$ | $D_{R3-R5}$ | $D_{R5-R6}$ | $D_{R6-R7}$ | $D_{R7-R8}$ | $T_{b_{SO-R1}}$ | $T_{b_{R1-R5}}$ | $T_{b_{R5-R7}}$ | $P_o$ | $P_s$ | $\Delta P_o$ | PBMA | Stacking |
| --- | --- | --- | --- | --- | --- | --- | --- | --- | --- | --- | --- | --- | --- | --- |
| m1 | 1/1 | 1/1 | 1/1 | 1/1 | 1/1 | 1/1 | 1/1 | 1/1 | 1/1 | 1/1 | 1/1 | 1/1 | 0.056 | 0.087 |
| m2 | 1/1 | 1/1 | 1/1 | 1/1 | 0/1 | 0/1 | 0/1 | 0/1 | 0/1 | 1/1 | 0/1 | 0/1 | 0.203 | 0.188 |
| m3 | 1/1 | 1/1 | 1/1 | 1/1 | 0/1 | 0/1 | 0/0 | 0/0 | 0/0 | 1/1 | 0/1 | 0/1 | 0.159 | 0.009 |
| m4 | 1/1 | 1/1 | 1/1 | 0/1 | 0/1 | 0/1 | 0/0 | 0/0 | 0/0 | 1/1 | 0/1 | 0/1 | 0.266 | 0.280 |
| m5 | 1/1 | 1/1 | 0/1 | 0/1 | 0/1 | 0/1 | 0/0 | 0/0 | 0/0 | 1/1 | 0/1 | 0/1 | 0.196 | 0.147 |
| m6 | 1/1 | 1/1 | 1/1 | 1/1 | 0/1 | 0/0 | 0/0 | 0/0 | 0/0 | 1/1 | 0/1 | 0/0 | 0.001 | 0.062 |
| m7 | 1/1 | 1/1 | 1/1 | 1/1 | 0/1 | 0/0 | 0/0 | 0/0 | 0/0 | 1/1 | 0/1 | 0/1 | 0.023 | 0.080 |
| m8 | 1/1 | 1/1 | 1/0 | 1/1 | 0/1 | 0/1 | 0/0 | 0/0 | 0/0 | 1/1 | 0/1 | 0/1 | 0.096 | 0.146 |
| m9 | 1/1 | 1/1 | 1/1 | 1/1 | 0/0 | 0/1 | 0/0 | 0/0 | 0/0 | 1/1 | 0/1 | 0/1 | 0.000 | 0.000 |

### 3.4 Developmental drivers of soybean maturity groups

Optimum photoperiod *P*_o_, and photothermal days *D*_R0-R1_, *D*_R1-R3_ and *D*_R3-R5_, were linearly related to MG (Figure 5, a). Photothermal days *D*_R5-R6_, *D*_R6-R7_, calendar days *D*_R7-R8_, photoperiod sensitivity *P*_s_ and the reduction in critical photoperiod threshold after R1,Δ*P*_o_, were unrelated to MG but differed between genotypes (Figure 5, b). Base temperatures 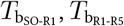and 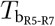were constant across genotypes and therefore can be considered species-specific (Figure 5, c). Optimum photoperiod *P*_o_ was the parameter most closely related to MG (Figure 5, a ). Despite 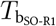(0.054 °C, CI: -0.03 to 0.14 °C) was slightly higher than 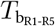(-0.07 °C, CI: -0.15 to 0.013 °C, probability of direction [PD] = 96%), this difference was almost negligible when calculating the temperature multiplier *ϕ*_T_ for stages SO-R1 and R5-R7 (Figure 5, d). Base temperature for grain filling 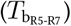was negative (-19.2 °C, CI: -19.8 to -18.5 °C), much lower than vegetative 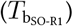and post-flowering 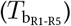base temperatures (Figure 5, c). This is in spite soybean does not survive freezing temperatures. However, this negative base temperature enhanced the predictive ability of our model, similarly to the findings of Boote et al. (1998b). The photoperiod multiplier *ϕ*_P_ as a function of photoperiod (Figure 5, e) showed a marked difference in *P*_o_ between contrasting genotypes DM 2200 (MG 2, 13.0 h, CI: 12.9 to 13.0 h) and RA 550 (MG 5.5, 12.4 h, CI: 12.4 to 12.5 h), but the slopes in the photoperiod response region were not visually evident in spite their differences in *P*_s_ (0.292 h^-1^, CI: 0.289 to 0.295 h ^-1^ vs 0.296 h ^-1^, CI: 0.293 to 0.299 h ^-1^, PD = 92%)

**Figure 5:**
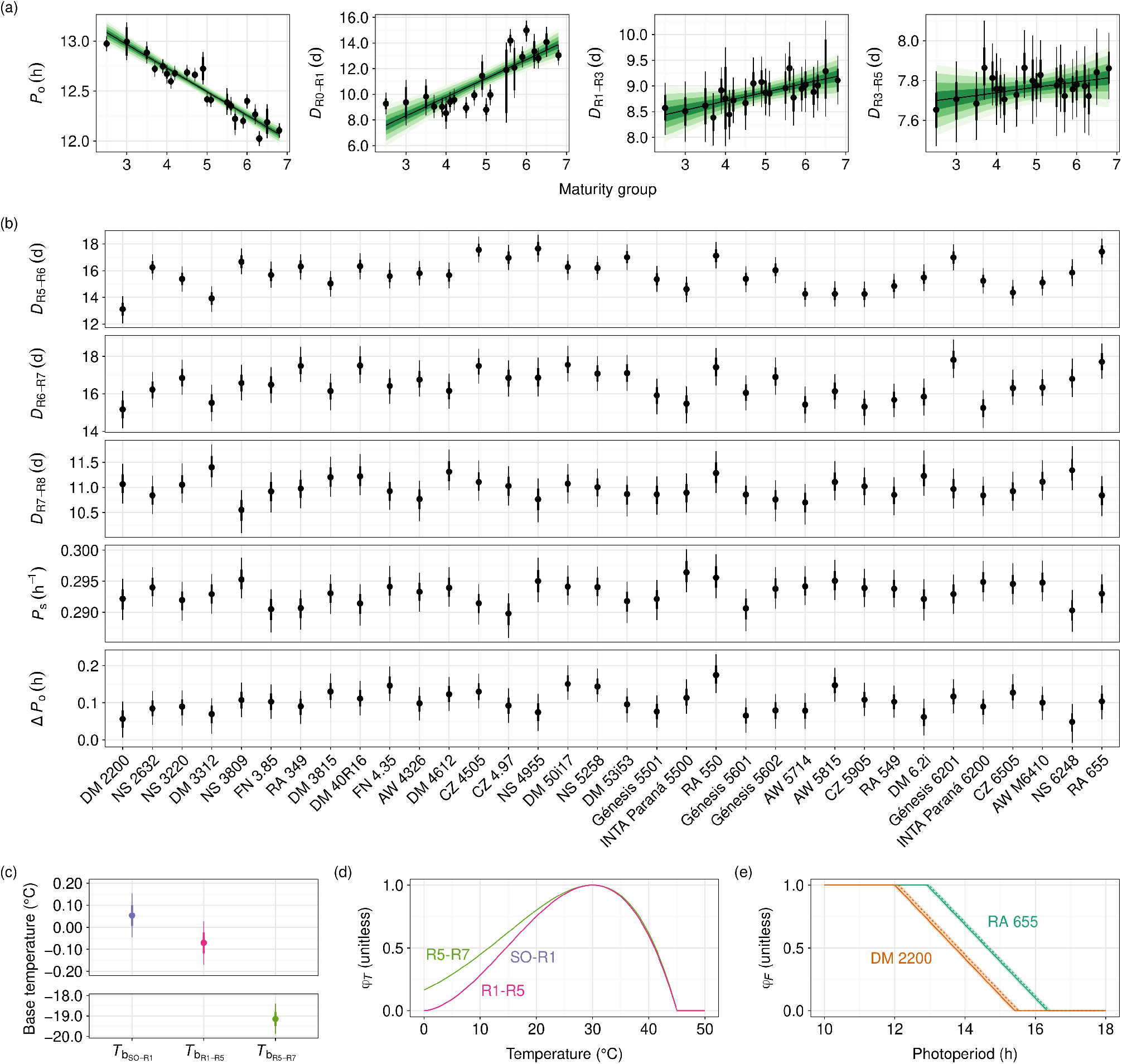
Parameters of model `u4` (Table 5) as modelled by the multivariate model `m4` (Table 6). (a) Parameters *P*_o_, *D*_R0-R1_, *D*_R1-R3_ and *D*_R3-R5_ as a function of maturity group, where the solid line represents the median regression line and green shades depict credible intervals 50, 90, 95 and 99%. (b) Parameters that showed no relationship with maturity group but a random effect (random intercept) of genotype instead. (c) Genotype-independent base temperatures 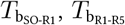and 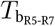. (d) Temperature multiplier *ϕ*_*T*_ as a function of temperature for stages sowing-R1 (SO-R1) R1-R5 and R5-R7. Notice that curves for SO-R1 and R1-R5 overlap each other almost entirely and thus cannot be visually distinguished (despite their different base temperatures). (e) Photoperiod multiplier *ϕ*_*P*_ as a function of photoperiod for the most contrasting genotypes in terms of *P*_o_, *P*_s_ andΔ*P*_o_ (DM 2200 [MG 2] and RA 550 [MG 5.5]), and for stages before (solid line) and after R1 (dashed line). From (a) to (e), points show the median fitted value and bars represent the 50 (thicker) and 90% (thinner) credible intervals.

### 3.5 Model’s ability to predict phenology

To assess the phenological accuracy of the calibrated model, each genotype’s prediction was validated against the validation data set. The model accurately predicted soybean phenology with an increased prediction error at later phenological stages (Table 7). The average RMSE for stages R1, R3, R5, R6, R7 and R8 was 5.8, 7.1, 7.4, 9.4, 8.5, and 9.5 days, respectively, representing 12.1, 11.0, 9.4, 9.3, 7.3, 7.5% in terms of the mean observed days after sowing (DAS) of each stage (Table 7). From those errors, the RLA, measuring the percentage of error given by a systematic deviation from the 1:1 line, was 43.1, 5.3, 15.1, 8.8, 17.2, and 17.2% for stages R1, R3, R5, R6, R7 and R8, respectively (Table 7), showing that this deviation was much higher for R1 than the rest of the stages. On a location basis, the RMSE was highest for Manfredi (Table 7), but these errors were less biased than those of Salta (17.2 vs 83.2% in RLA; Figure 6 and Figure 7). Salta showed biased errors towards the end of the reproductive stage (R6, R7 and R8, Table 7, Figure 6 and Figure 7). Overall, the model showed a very good level of prediction with RMSE values from 5.8 to 9.5 days and RRMSE from 12.1 to 7.5% for R1 to R8 developmental stages.

**Table 7:** The ability of the multivariate model to predict developmental stages across all genotypes. Root mean squared errors (RMSE, days), relative root mean squared errors (RRMSE, %), and relative lack of accuracy (RLA, %) for all tested locations as well as across them (total) for stages R1, R3, R5, R6, R7 and R8.

|  | Metric | Stage |  |  |  |  |  | Total |
| --- | --- | --- | --- | --- | --- | --- | --- | --- |
|  |  | R1 | R3 | R5 | R6 | R7 | R8 |  |
| Capitán Miranda | RMSE (d) | 3.5 | 4.9 | 3.9 | 5.6 | 5.5 | 6.4 | 5.1 |
|  | RRMSE (%) | 9.1 | 8.7 | 6.0 | 6.5 | 5.5 | 5.9 | 6.7 |
|  | RLA (%) | 18.2 | 55.9 | 26.6 | 44.7 | 42.3 | 50.5 | 11.6 |
| Chascomús | RMSE (d) | 7.3 | 4.1 | 6.3 | — | 3.5 | 4.6 | 5.9 |
|  | RRMSE (%) | 11.4 | 5.4 | 7.2 | — | 2.5 | 2.9 | 7.2 |
|  | RLA (%) | 74.6 | 39.1 | 58.1 | — | 100.0 | 100.0 | 2.1 |
| La Estanzuela | RMSE (d) | 6.0 | 6.4 | 5.6 | 10.4 | 6.2 | 8.8 | 7.5 |
|  | RRMSE (%) | 10.8 | 9.4 | 6.8 | 9.4 | 5.0 | 6.6 | 7.8 |
|  | RLA (%) | 39.8 | 10.5 | 1.3 | 54.2 | 34.0 | 32.6 | 2.1 |
| Manfredi | RMSE (d) | 7.1 | 10.7 | 10.6 | 15.6 | 11.4 | 11.2 | 11.3 |
|  | RRMSE (%) | 14.9 | 15.9 | 13.0 | 14.2 | 9.1 | 8.3 | 12.3 |
|  | RLA (%) | 30.3 | 22.8 | 30.7 | 43.6 | 9.5 | 3.5 | 17.1 |
| Pergamino | RMSE (d) | 7.5 | 7.8 | 8.6 | 7.3 | 6.5 | 9.0 | 7.8 |
|  | RRMSE (%) | 13.7 | 10.9 | 9.5 | 6.7 | 5.2 | 7.0 | 8.3 |
|  | RLA (%) | 59.2 | 42.1 | 54.9 | 15.8 | 53.8 | 46.9 | 2.2 |
| Reconquista | RMSE (d) | 4.1 | 4.2 | 6.3 | 3.9 | 10.0 | 8.4 | 6.1 |
|  | RRMSE (%) | 10.7 | 7.6 | 9.2 | 4.6 | 9.7 | 6.9 | 8.7 |
|  | RLA (%) | 36.8 | 30.8 | 72.0 | 22.3 | 17.1 | 11.3 | 0.8 |
| Salta | RMSE (d) | 2.0 | 6.5 | 5.5 | 11.6 | 18.6 | 19.6 | 9.8 |
|  | RRMSE (%) | 4.5 | 11.8 | 8.0 | 12.4 | 16.8 | 16.1 | 14.0 |
|  | RLA (%) | 4.7 | 81.4 | 88.2 | 84.9 | 96.0 | 96.2 | 82.4 |
| Total | RMSE (d) | 5.8 | 7.1 | 7.4 | 9.4 | 8.6 | 9.6 | 7.9 |
|  | RRMSE (%) | 12.0 | 11.1 | 9.4 | 9.3 | 7.3 | 7.6 | 9.3 |
|  | RLA (%) | 42.8 | 5.3 | 15.1 | 9.2 | 17.0 | 17.3 | 0.7 |

**Figure 6:**
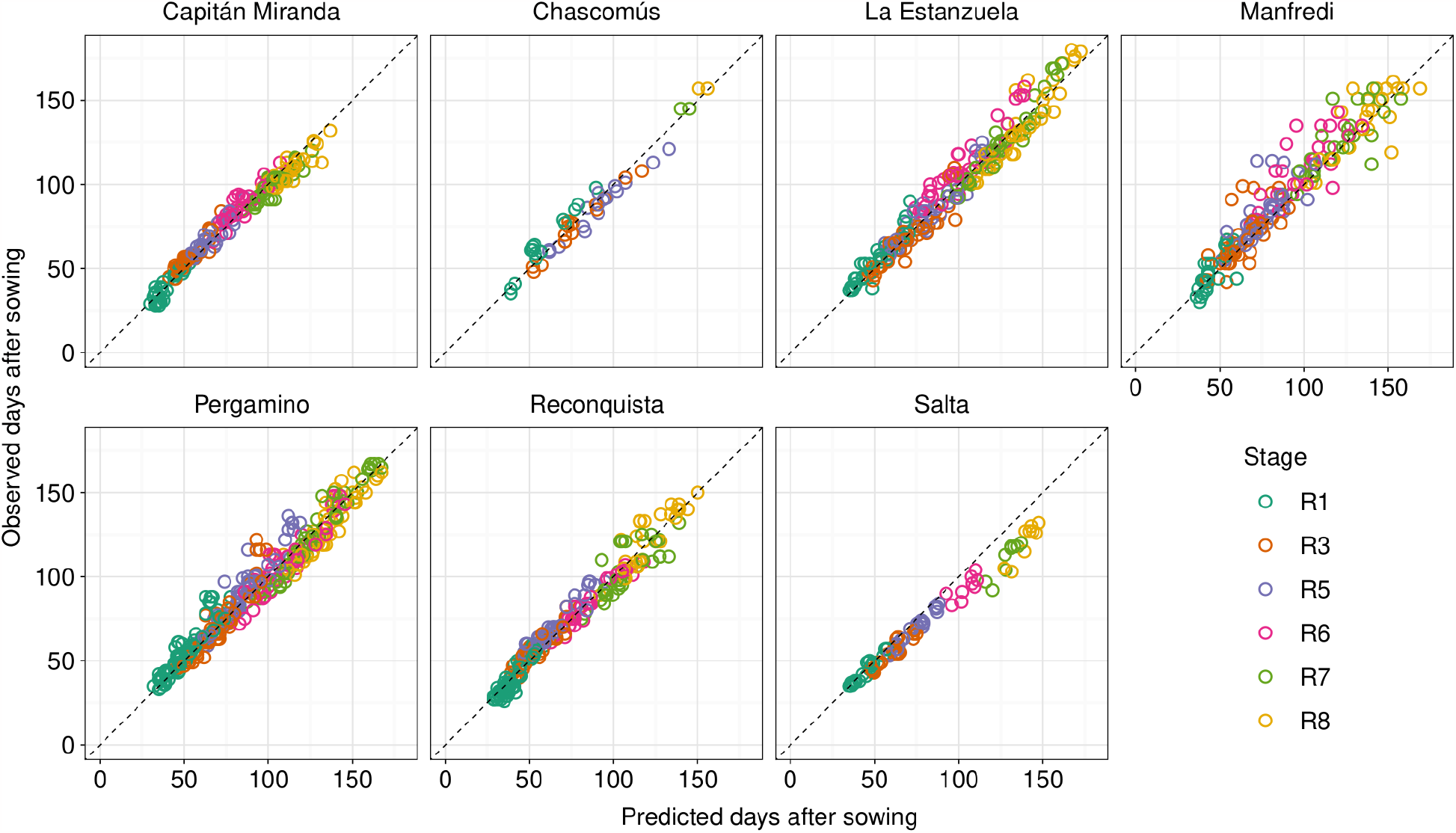
Observed days after sowing as a function of predicted days after sowing for stages R1, R3, R5, R6, R7 and R8 evaluated with data from the validation data sets at all sites using the multivariate model. The dashed line represents the *y* = *x* line.

**Figure 7:**
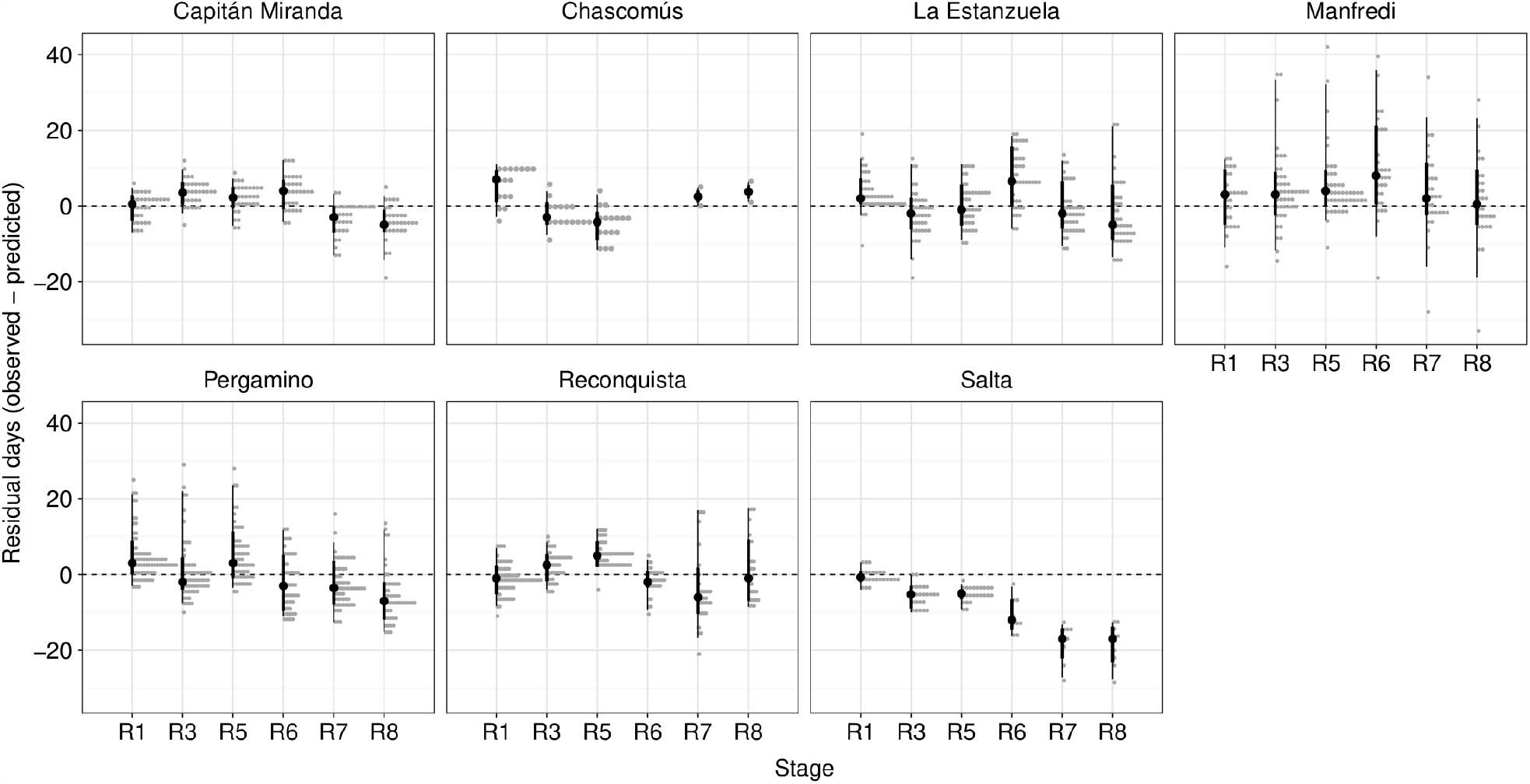
Evaluation of the difference between observed and predicted days after sowing for stages R1, R3, R5, R6, R7 and R8 at all sites using the validation data set. Black points represent the median, whereas the thick and thinner lines show the 50 and 90% credible intervals. Grey dots show the distribution of the residuals as counts

### 3.6 Model predictions for website deployment

The algorithms generated to predict each of the developmental stages of the soybean crop were included in a software named CRONOSOJA (http://cronosoja.agro.uba.ar/). The software is simple and friendly for the users, who can select a location within the country (Argentina, Paraguay or Uruguay), a genotype, and a particular sowing date. The model returns predicted dates of the different phenological stages with an estimated error value. In addition, the CRONOSOJA model indicates the critical period of the crop, the risk of frost for each of the phenological stages and the dynamics of plant available water in the profile (Figure 8).

**Figure 8:**
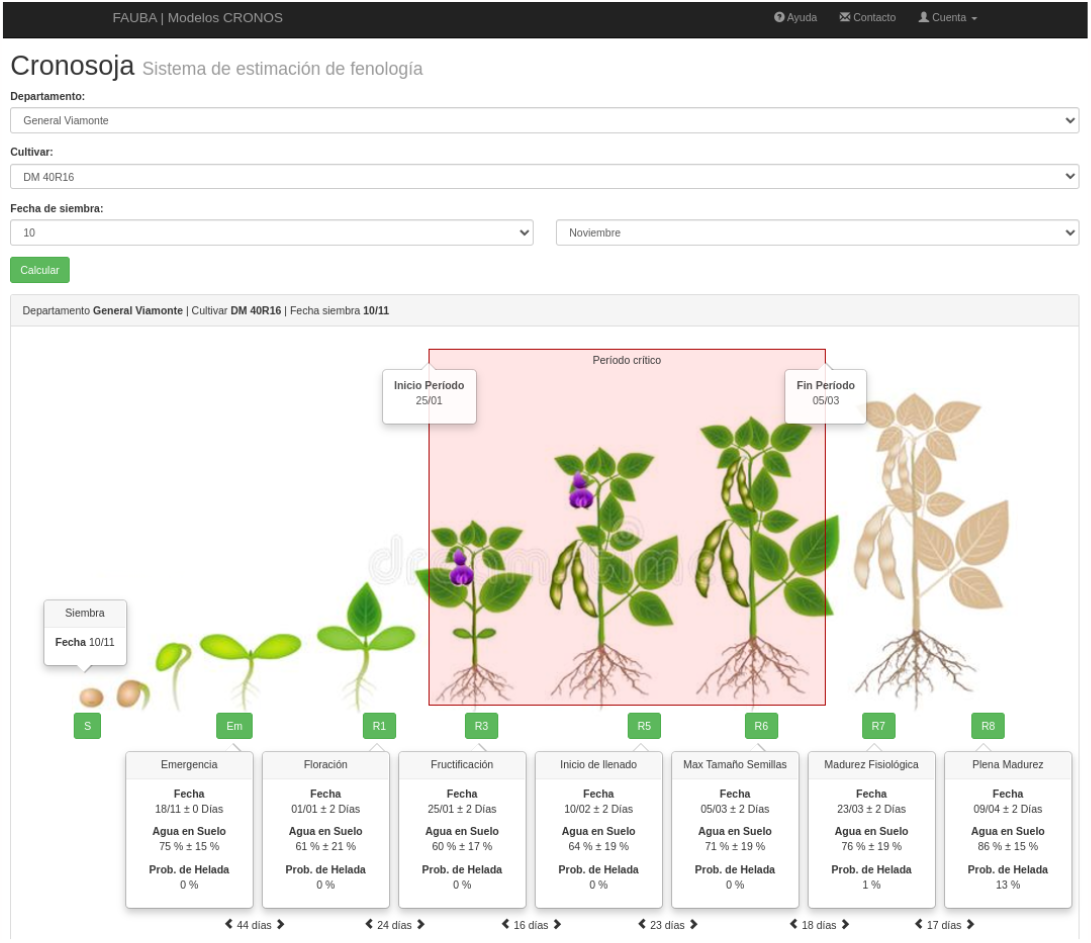
Example of the output information from the CRONOSOJA model in the web version where the prediction dates of the model for the different phenological stages are detailed. The red shaded rectangle indicates the time interval of occurrence of the critical period.

## 4 Discussion

Field experiments in different countries (Argentina, Uruguay and Paraguay) were conducted to evaluate phenology in soybean varieties, grown over a range of latitudes from -24.9 to -35.6, and at different sowing dates between October and January, allowing us to explore a wide range of thermo-photoperiodic conditions. The combination of a wide range of environmental conditions and MG from 2 to 6 allowed us to precisely dissect soybean developmental responses to the environment. In this context, three base temperatures that differed across developmental stages and an extra parameter for calculating the photoperiod multiplier after R1 were identified, in line with previous studies (Boote et al., 2003; Grimm et al., 1994; Grimm et al., 1993; Piper et al., 1996). A wide range of variation in the length of the phases from sowing to different developmental stages across different genotypes was explored to select between alternative phenological models that best explained the observed experimental data.

A rapid method of model selection by means of best predictive accuracy was employed (Vehtari et al., 2017) in a dynamic developmental model (Jones et al., 2003; Keating et al., 2003). This allowed us to rapidly assess which model structure was the best. An alternative would be to perform another type of cross-validation called k-fold, as used previously by Wallach et al. (2001), but this would have been unfeasible in our case as it would have taken much more computational time given the large number of iterations required to reach convergence of the Bayesian model. Using our framework as a proxy for model parsimony, we were able to choose meaningful structures for our models, whereby our understanding of the processes determining phenology was increased. For instance, we were able to corroborate in field conditions that soybean’s sensitivity to photoperiod increased after flowering (Table 5). The parameter representing this extra sensitivity,Δ*P*_o_, is already included in DSSAT-CROPGRO as a parameter called R1PPO (Boote et al., 1998a) but is absent in APSIM Soybean (Robertson et al., 2002). Regarding base temperature, some models suggest a constant value for predicting soybean phenology. Cober et al. (2001) proposed a constant base temperature of 5.8 °C to predict flowering time. Similarly, Hadley et al. (1984) and Major et al. (1975a) showed that base temperature to predict flowering time, when photoperiods were longer than the optimum value, was 7.8 °C without differences among the used varieties. However, a model with variable base temperatures, during vegetative, reproductive and grain filling phases, was better than a model considering constant base temperatures (Table 5). Moreover, we found lower base temperatures compared to those authors, although this may obey to our choice of a curvilinear function instead of a linear one to explain temperature effects (Equation 2, Equation 3, Figure 5, d). Variable base temperatures are already formalised in DSSAT-CROPGRO (Boote et al., 1998b; Grimm et al., 1993) but not in APSIM Soybean, which uses a constant base temperature (Robertson et al., 2002). The present work suggests that the latter modelling platform could benefit from includingΔ*P*_o_ and a stage-dependent *T*_b_ to its inner functions.

Model selection also showed that changes in the determinants of phenology across maturity groups were mainly due to processes occurring early in reproductive development, around phases EM-R1, R1-R3 and R3-R5 (Figure 5, a). Moreover, the differences between cultivars were due to photoperiod sensitivity, with similar responses to temperature (Grimm et al., 1993). In contrast to Piper et al. (1996), who showed no association between parameters controlling the duration of R1-R5 phases, we discovered that photothermal days for these stages were a linear function of maturity group. However, *P*_o_, the optimum photoperiod affecting all developmental stages between the end of the juvenile phase and physiological maturity, was the parameter that best correlated with the maturity group. This finding suggests that *P*_o_ may be a good proxy to estimate maturity groups when there are ambiguities in their determination by seed companies. We did not detect a relationship between maturity group and photoperiod sensitivity, *P*_s_, as reported by Piper et al. (1996) and as it is parameterised in DSSAT-CROPGRO model (Boote et al., 2003). This may be due to a weakness of model fitting given the high of collinearity between *P*_o_ and *P*_s_, in spite our attempt to improve this issue by imposing mild informative priors over them (Table 4).

The ability of the model to predict phenology was greater for early stages of development (Table 7). Predictions for later stages were less accurate, in accordance with Salmerón and Purcell (2016). This is in agreement with our pre-modelling quantification of sources of variation for stages duration, which showed that the fraction of unexplained variance increased towards the end of the developmental cycle (Figure 4). The model underpredicted development rate in cool environments like Salta, an issue already reported by Piper et al. (1996) that may highlight a weakness of the model to represent soybean development at low temperatures.

The outputs of the selected model that works at the level of genotypes or maturity group were included within a user-friendly interphase called CRONOSOJA (http://cronosoja.agro.uba.ar/), which is available for free and includes all soybean productive regions from Argentina, Paraguay and Uruguay.

## 5 Conclusions

The results of the study showed that parameters concerning early plant development, such as those affecting stages of vegetative growth and flowering, were the most affected by the different maturity groups (MG). However, base temperatures did not change across MG or between genotypes. The optimum photoperiod *P*_o_, and photothermal days *D*_R0-R1_, *D*_R1-R3_ and *D*_R3-R5_, were linearly related to MG. Photothermal days *D*_R5-R6_, *D*_R6-R7_, calendar days *D*_R7-R8_, photoperiod sensitivity *P*_s_ and the reduction in optimum photoperiod threshold after R1,Δ*P*_o_, were unrelated to MG but differed among genotypes. Base temperatures 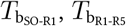and 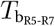, were constant across genotypes and can be considered species-specific. The optimum photoperiod *P*_o_ was the parameter most closely related to MG.

## 7 Acknowledgements

We thank Francisco Raspa, Luis Blanco, and Octavio Ghio Trebino for their dedication and hard work in conducting the field experiments, which greatly enhanced the quality and impact of this scientific article.

## Supporting information

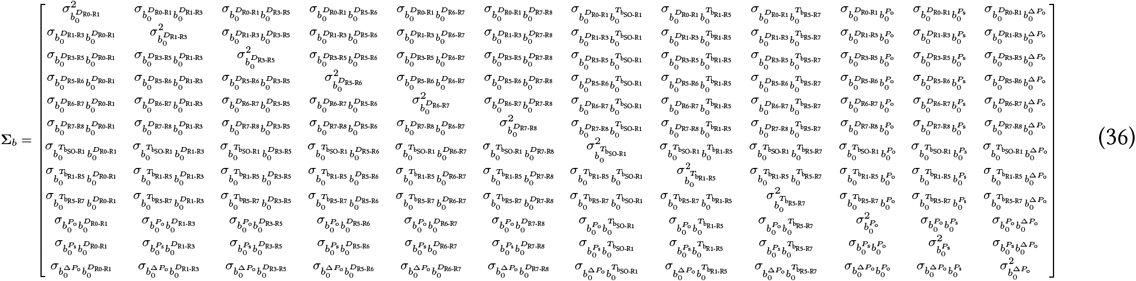

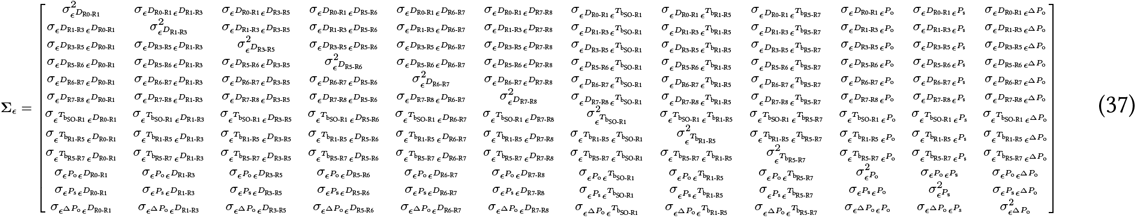

